# An automated workflow for the discovery and docking simulation of the protein-protein complexes using *in vivo* chemical cross-linking

**DOI:** 10.1101/373829

**Authors:** Vladimir Svetlov, Evgeny Nudler

## Introduction

Chemical cross(X)-link mapping assisted by mass spectrometry (XL-MS, also CXMS and CLMS) is a low-resolution hybrid method of structural biology, yielding a set of pairwise distance restraints between reactive solvent-accessible amino acids^1-9^. Most commonly used X-linkers at present belong to the class of amino-reactive homo-bifunctional NHS-esters, which act as protein proximity sensors, connecting predominantly Lys residues (as well as far less common and informative N-terminal amines)^10-14^. Main structural application of XL-MS to date is limited to the generation of distance restraints assisting in interpretation of the cryo-EM data^3,5,15-17^. This approach usually entails treating the same sample used in cryo-EM experiments (purified protein-protein complex (PPC) often assembled from over-expressed components) *in vitro* with chemical X-linkers prior to the entry into the XL-MS pipeline. In that XL-MS relies on an orthogonal information regarding PPC composition as do other structural methods, thereby limiting its exploratory potential.

Chemical X-linking has been used in proteomics and interactomics applications long before the advent of XL-MS. Most commonly *in vivo* interactomics approach utilizes small, broadly specific, cell-permeable X-linkers such as formaldehyde, administered to the culture/or suspension of cells (live tissue applications face an obvious impediment to X-linker diffusion due to multi-layering of membranes and cell walls, as well as intra-cellular macromolecular crowding)^18-21^. X-linked PPCs of interest are then purified, using affinity (including immuno-affinity) tags, and subjected to proteolysis followed by MS/MS identification of linear peptides(AP/IP-MS)^22-24^. This enumerative proteomics is made possible by the fact that X-linkers react only with solvent-accessible amino acids, leaving buried/interior peptides unmodified and thus amenable to straightforward shotgun protein ID.

Non-labile covalent X-links formed *in vivo* antagonize dissociation of the PPC during cell disruption, extraction and purification steps, thereby reducing generation of the false negative data points compared to affinity/immune-purification carried out without X-linking. However, lacking means to distinguish co-purification from specific PPC formation, it does not prevent inclusion of the false positive points into the dataset, which often includes hundreds of potential interactors for any given “bait” (affinity-tagged target protein). In contrast, XL-MS by providing objective measure of proximity and unique PPC architecture (as a non-random set of X-linked peptides) allows for an unbiased removal of the false positives from the data.

XL-MS use in *in vivo* interactomics has been rather limited. Formaldehyde, although widely used in X-linking applications, has broad reactivity towards amino acids, thereby complicating automated (rule-based) discovery^21^. In contrast, popular XL-MS cross-linkers often exhibit low cell-permeability, due to the presence of easily ionizable groups (e.g. bis(sulfosuccinimidyl)suberate (BS3)) or bulky spacers (e.g. functionalized reagents featuring affinity or clickable groups). Successful forays into XL-MS of live cells have been rather scarce, with one of the most successful platforms developed for *in vivo* XL-MS utilizes cleavable NHS-ester X-linker disuccinimidyl sulfoxide (DSSO) and LC-MS^n^-assisted identification of the X-linked peptides^25^. This platform offers many advantages to a highly skilled operator, but its reliance on in-house expertise and scripts certainly restricts its portability to the labs that do not have access to either. In this work we set to assemble a streamlined *in vivo* XL-MS workflow, from chemical X-linking to X-link-assisted protein-protein docking, which utilizes broadly available reagents and automated (score-driven) computational tools and thus requires minimal expert intervention.

## Results

### Discovery of *E. coli* NusA-RNA polymerase (RNAP) interactions by *in vivo* XL-MS

Water soluble X-linker BS3 has been successfully used in structural interrogation of PPCs by XL-MS *in vitro* due to its preference for Lys as the reactive group, predictable fragmentation of X-linked peptides, and commercial availability^16,17,26-29^. Its cell-permeable isostere disuccinimidyl suberate (DSS) was routinely deployed in AP/IP-MS proteomics experiments, as well as in *in vitro* XL-MS applications^30-35^. To extend its use to *in vivo* XL-MS we have treated an exponentially growing culture of *E. coli* RL721 (*rpoC::His6*) with 2 mM DSS for 30 min at 37°C, RNAP polymerase and the associated proteins were purified using Ni^2+^-affinity resin, and digested with trypsin for LC-MS/MS analysis. Each sample was divided in two aliquots, the first were processed for linear peptide discovery (data-dependent acquisition of ions charged 1, 2, and 3), and the second – for X-linked peptides discovery (data-dependent acquisition of ions charged 4, 5, and 6).

In order to reduce non-specific binding of proteins to the resin purification was carried out in denaturing conditions (in presence of 8M urea). The number of co-purifying proteins identified in the denatured samples was reduced compared to the same discovered in the samples purified without the addition of urea, although not entirely free from known contaminants and other unrelated proteins (data not shown). As expected from the His_6_-tagged RpoC-baited samples, the list of co-purified proteins included RpoC, other RNAP subunits (RpoB, RpoA, RpoZ), and RNAP-associated initiation and elongation factors (RpoD, RpoS, NusA, NusG, GreA, GreB, DksA, etc), as well as unrelated proteins of *E. coli* and human origins (VS and EN, manuscript in preparation). *fasta* files of all corresponding proteins were extracted from the Uniprot and pooled together to form the search database (combined fasta file) for XL-MS. The reduction of the search space to the proteins actually present in the sample lowers the computational cost (compared to the search of the entire proteome), but lacking any other filtering of the input, it prevents biasing the discovery process.

XL-MS results obtained by pLink1 were filtered during the search and recording of the output by the false discovery rate (FDR<5%) and e-value (≤0.001)^36^. As expected from the RpoC-baited AP-XL-MS, discovered X-links originated predominantly in one of the RNAP subunits (VS and EN, manuscript in preparation). No X-links between RpoA/B/C/Z proteins and known contaminants (e.g. metal-binding metabolic enzymes)^37^ were detected in the sample, indicating that this approach didn’t introduce any false positives into the dataset. Consistent with orthogonal interactomics data, a number X-links were detected between RNAP subunits and potential interactors, such as initiation and elongation factors RpoD, Rho, DksA, NusG, and NusA. Efficient discovery of these X-links in a crude, single-tag AP is most likely due to the high abundance of these factors, comparable with that of RpoC (7164 molecules/cell in LB): RpoD - 1657, Rho – 5934, DksA – 19594, NusG – 6938, and NusA – 10025 molecules/cell, respectively^38^. It bears noting that no X-links were found between RNAP subunits and highly abundant unlikely interactors, such as SucB (15565), GltA (16361), and CysK (21811, all – molecules/cell). The apparent lack of the experimental X-links between Rpo proteins and low abundance transcriptional regulators present in the sample, such as MalT (75 molecules/cell) and ExuR (94 molecules/cell), indicates that this approach is prone to generate false negatives in regard to under-represented proteins, when highly abundant bait is used. This predicament may be endemic to the highly complex RNAP interactome, which at any given time consists of hundreds of transcription regulators present at intracellular concentrations spanning 4 orders of magnitude^39-41^.

For the purpose of generating an automated PPC docking model based on *in vivo* XL-MS data we selected the binary interaction between *E. coli* RNAP and the general transcription elongation regulator NusA. NusA was found in the cell in the concentration nearly stoichiometric to that of RNAP subunits, and is believed to be present in the majority, if not the entirety of the actively transcribing elongation complexes^42,43^. NusA plays a variety of roles in transcript elongation and termination, as well as in a continually growing number of transcription-associated/-coupled cellular functions, pertaining to DNA repair and genome stability, protein traffic on DNA, etc^44-54^. The availability of orthogonal structural information regarding NusA-RNAP allows for a confident benchmarking of the docking model. In XL-MS experiments NusA-RNAP X-links were highly abundant and reproducible, consistent with the existence of a specific and persistent PPC.

Majority of the *in vivo* NusA-RNAP X-links exhibited clustering in the N-terminal domain (NTD) of NusA and the vicinity of the so-called beta flap domain of RNAP, with the remainder scattered around flexible regions in the C-terminal half of NusA and flexible/unstructured fragments of RpoA subunit(s) of RNAP. Hence we have selected the following set of *in vivo* X-links formed between NusA NTD and RNAP for the docking simulation: NusA3-RpoB900, NusA37-RpoB890, NusA37-RpoB909, NusA38-RpoB890, NusA38-RpoB909, NusA111-RpoB890, NusA111-RpoB909, and NusA143-RpoC50.

### X-links-guided docking of NusA NTD and RNAP

There are a few protein-protein docking algorithms/servers, with HADDOCK standing out as the one that performs consistently well in the automated mode, with the starting structural files and unambiguous distance restraints as the only input from the user^55^. Lacking a high quality experimental structure of the full-length NusA, we have modeled its NTD (residues 1-180) using one of the top performing automated homology modeling servers, I-TASSER^56^. Resulting high-confidence model (C-score=0.98, estimated TM-score=0.85±0.08, estimated RMSD=2.7±2.0 Å) was further refined using FG-MD server^57^ to yield NusA NTD starting structure file, chain A. Since the only experimental inter-protein X-links formed by NusA NTD were limited to the RNAP two largest subunits, RpoB and RpoC, we have extracted coordinates of those subunits from experimental X-ray crystallographic model 4lk1^58^. Chains corresponding to RpoB and RpoC were joined into one (chain B), numbering of the RpoB residues was left unaltered, whereas RpoC residues were re-numbered by adding 2000 to their original numbers.

Unambiguous restraints were entered as the pairwise NZ distances between residues NusA3-RpoB900, NusA37-RpoB890, NusA37-RpoB909, NusA38-RpoB890, NusA38-RpoB909, NusA111-RpoB890, NusA111-RpoB909, and NusA143-RpoC50, conservatively set at 10.0 Å as the target distance, 4.0 Å as the lower correction, and 3.0 Å as the upper correction^59,60^. Docking was performed using the Expert Interface at the HADDOCK server, yielding 200 docking models divided between 2 clusters. Cluster 1 comprised 117 models and had higher HADDOCK score (127.0±9.0), greater buried surface area (1901.2±134.6), and lower restraints violation penalty (87.9±6.99), than cluster 2 (for the detailed comparison see Figure 1 and Table 1). Given the focus of this work on the automated, score-driven workflow we have selected cluster 1 as the docking outcome based on its superior scores, as provided by HADDOCK. Top model (model 1) from this cluster was refined using HADDOCK refinement interface and used in subsequent evaluation (Fig 2A).

**Figure 1.**
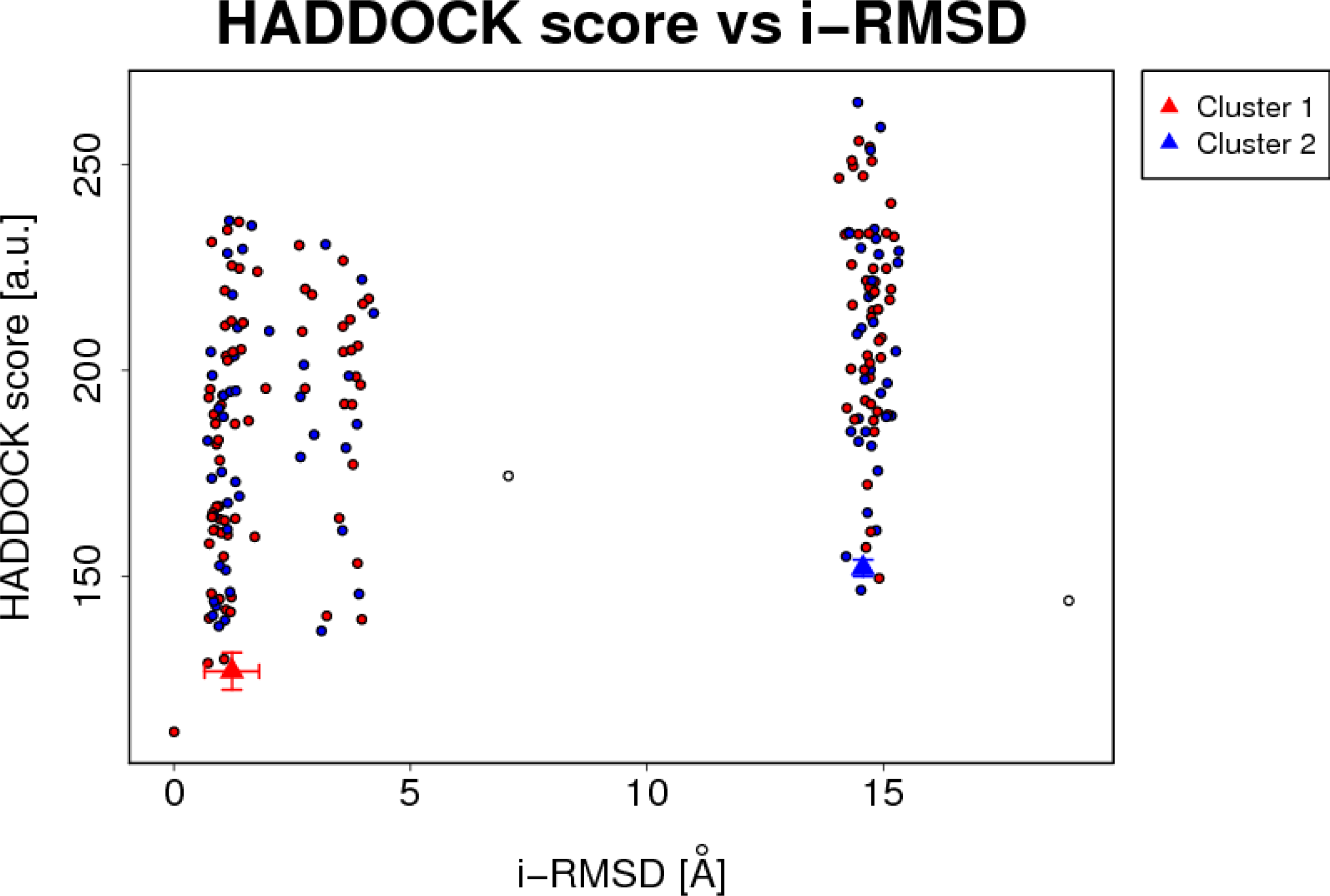
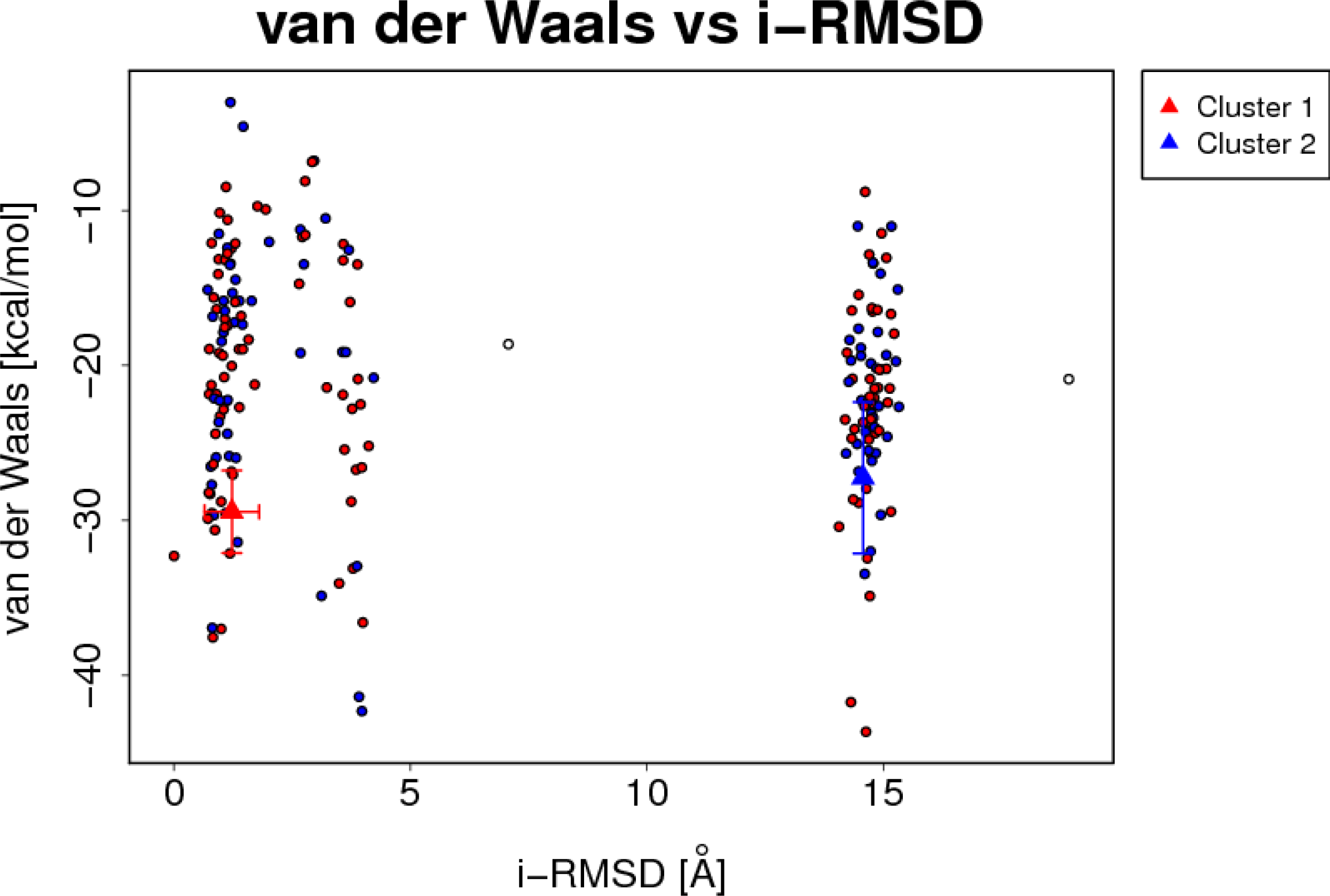
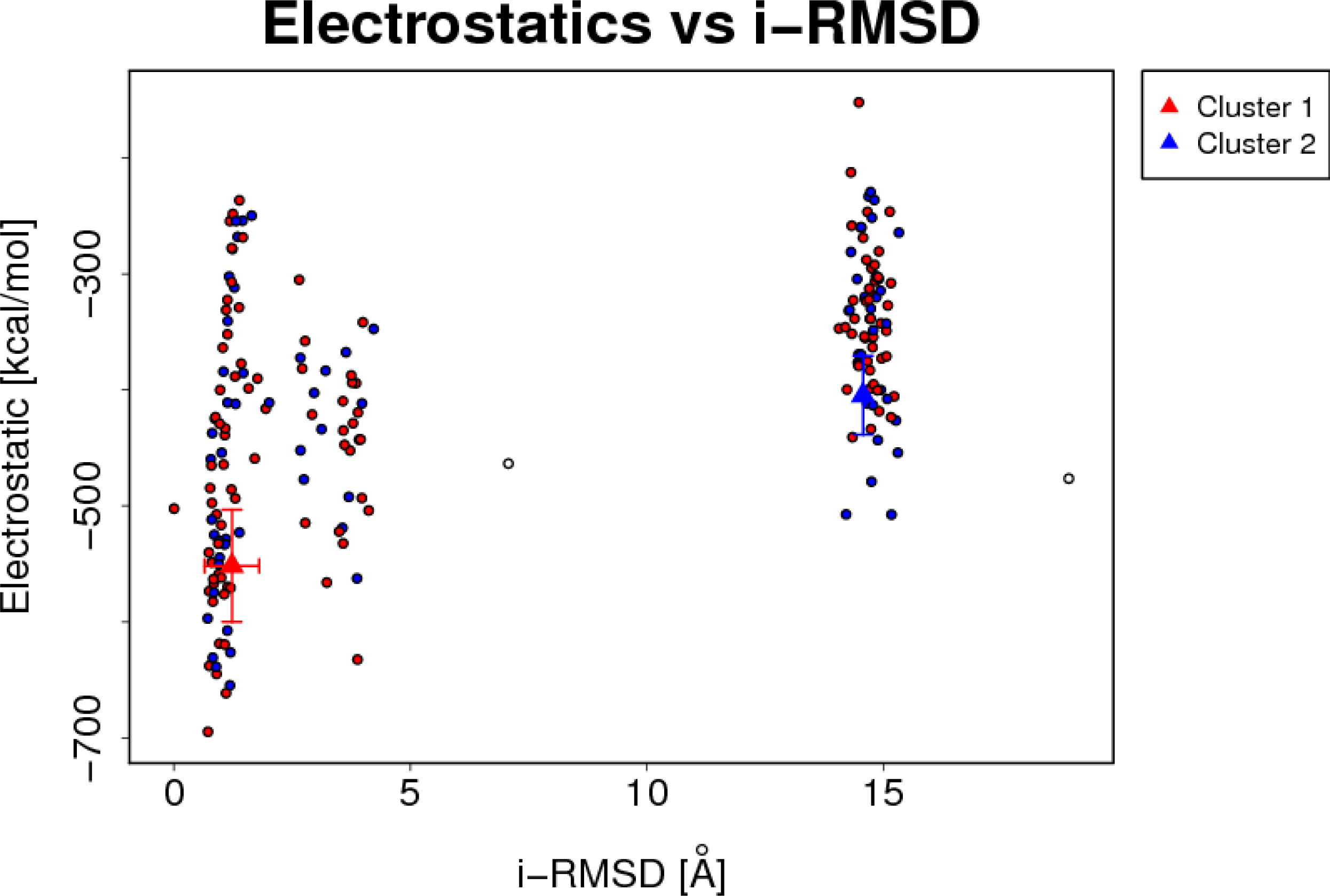
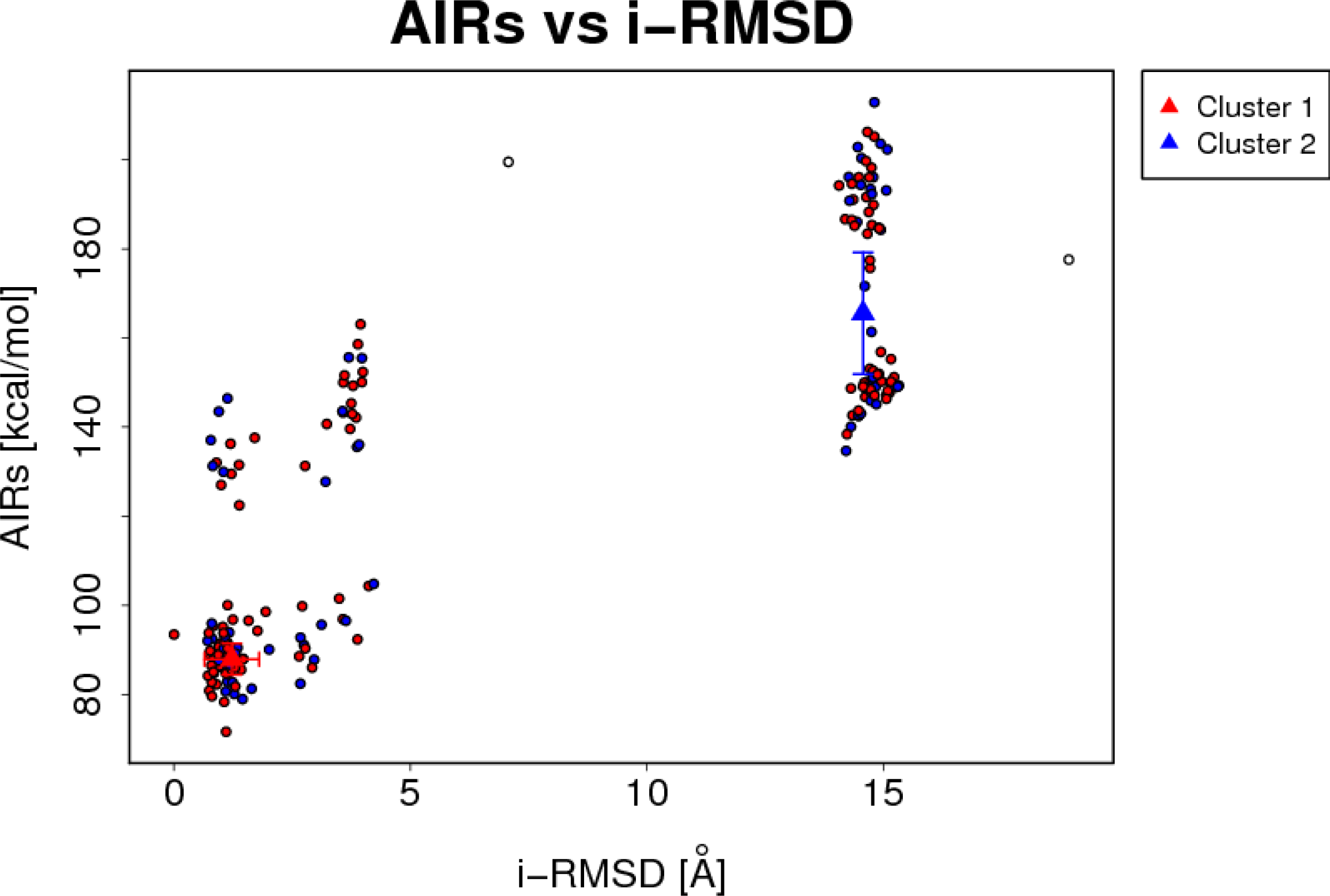
Scatter plot of the RMSD vs HADDOCK metrics for clusters 1 and 2. A. HADDOCK score; **B**: van der Waals energy; **C**: electrostatic energy, **D**: distance restraints violations (AIRs).

**Table 1.** Comparison of HADDOCK-generated metrics for NusA-NTD/RNAP models clusters 1 and 2

|  | Cluster 1 | Cluster 2 |
| --- | --- | --- |
| HADDOCK score | 127.0±9.0 | 152.0±4.1 |
| Cluster size | 117 | 81 |
| RMSD from the lowest-energy structure | 0.9±0.8 | 8.7±0.1 |
| Van der Waals energy | -29.5±5.3 | -27.2±9.8 |
| Electrostatic energy | -551.8±96.7 | -404.7±68.1 |
| Desolvation energy | 258.0±19.8 | 243.7±16.0 |
| Restraints violation energy | 87.9±6.99 | 165.5±27.32 |
| Buried surface area | 1901.2±134.6 | 1386.1±154.3 |
| Z-score | -1.0 | 1.0 |

**Figure 2.**
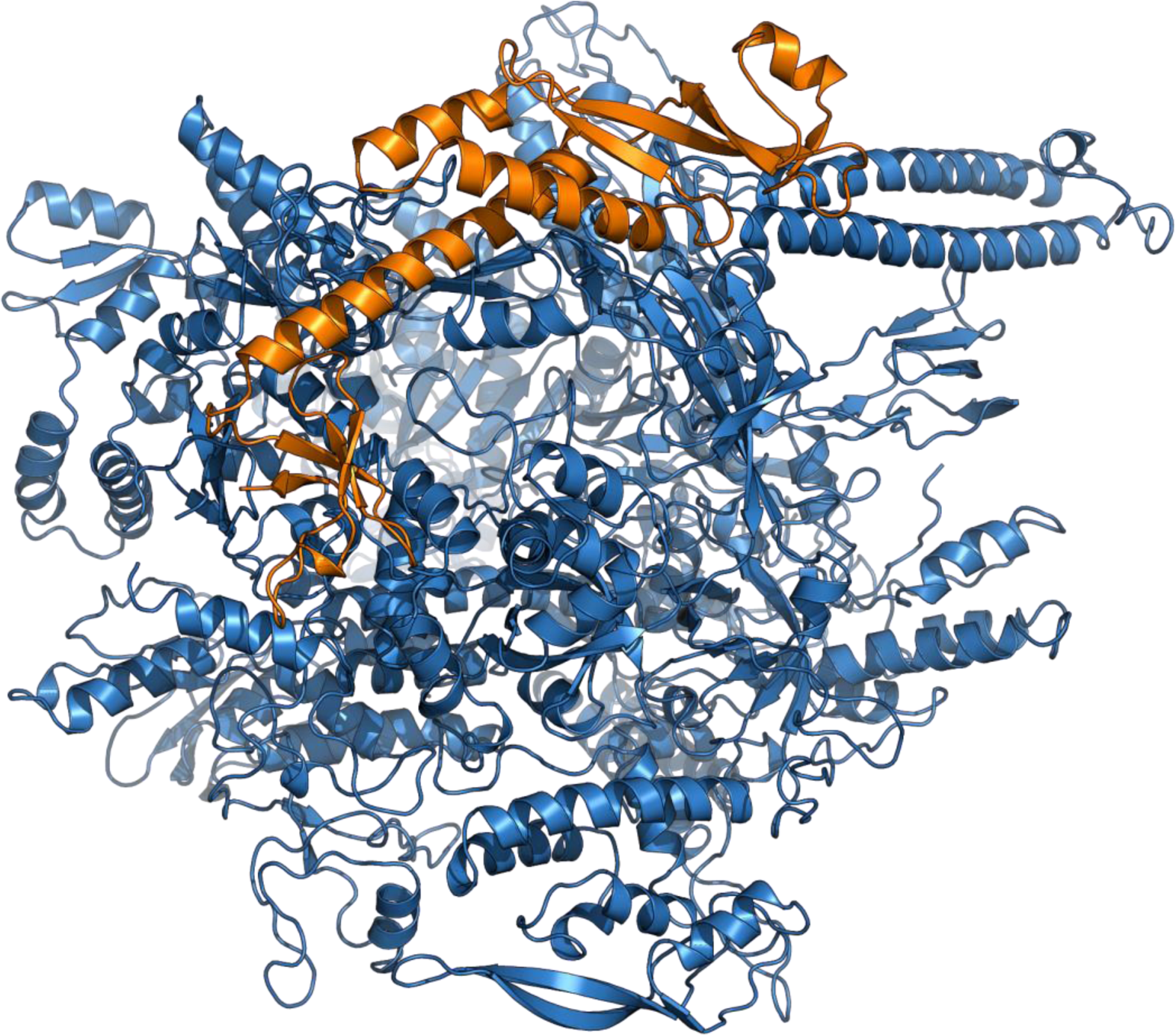
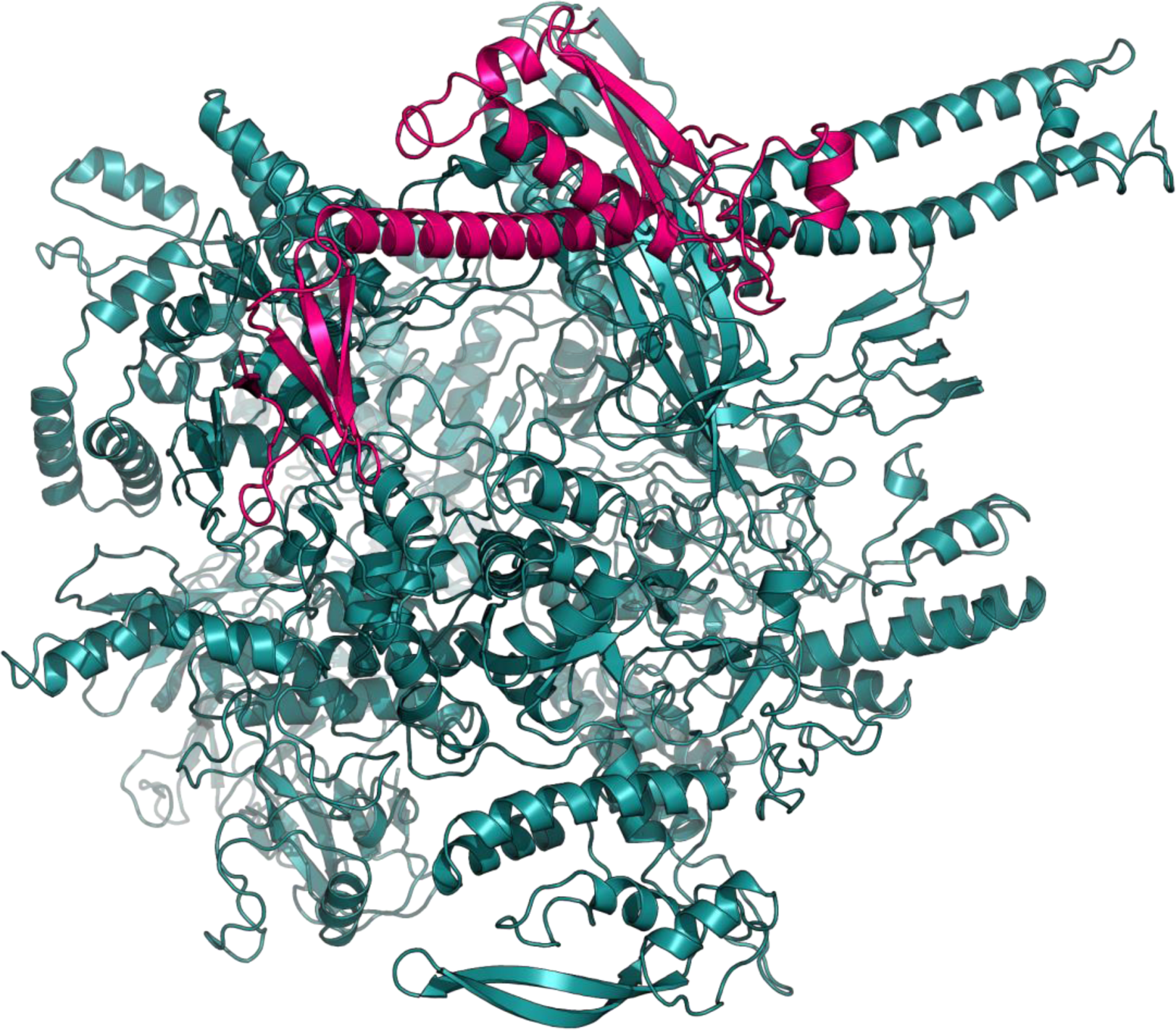
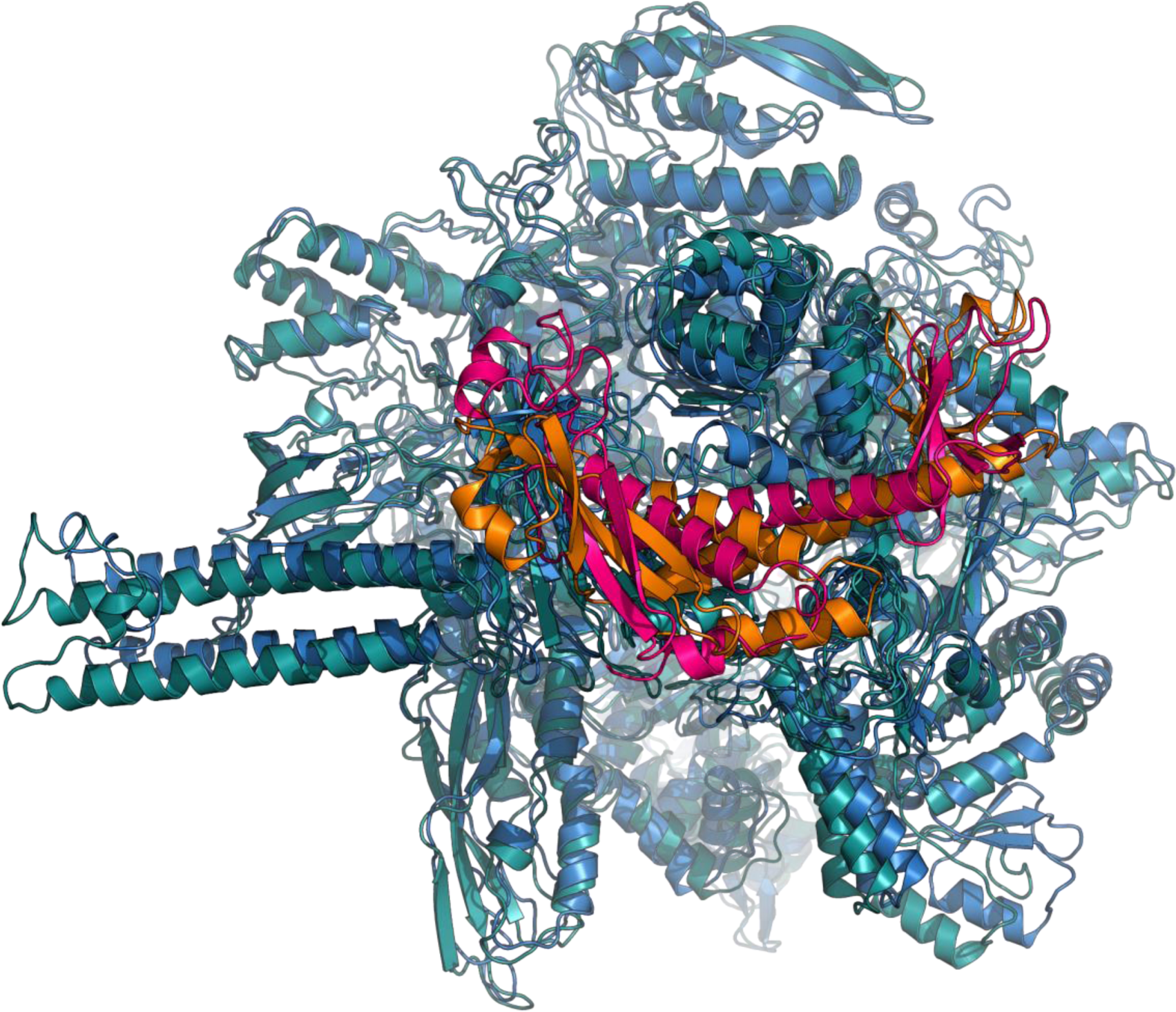
Structures of NusA-NTD/RNAP complex. **A**: docking model (RNAP=blue cartoon, NusA-NTD=orange cartoon); **B**: cryo EM model (RNAP=teal cartoon, NusA-NTD=hot pink cartoon); **C**: superimposed docking (A) and cryo EM(B) models.

### Comparison of the NusA NTD/RNAP docking and experimental structural models

Coordinates for the experimental model of *E. coli* NusA-NTD were extracted from the cryo electron microscopy structure deposited as 6flq^61^, namely the chains C, D, and F for RpoB, RpoC, and NusA, respectively. Chain F was trimmed to the first 180 residues to match the size of NusA NTD used in docking simulation (Fig 2B). Superimposition (*super* command in PyMOL^62^) revealed good overall congruity of these two models (Figure 2C), with NusA NTD binding in the same orientation to the same (beta flap) region of RNAP (consistent with other structural and biochemical data)^46,63-65^. TM-score^66^ of alignment, 0.4955, indicates near identity of the overall fold, and is very close to the TM-score of the aligned models without NusA NTD contribution (0.5450) (TM-scores of different experimental *E. coli* RNAP structure alignments vary between 0.5010 and 0.6205).

In order to ascertain the degree of agreement between the structural models (experimental and docking) and the experimental NusA NTD-RNAP X-links we have calculated the Euclidean CA-CA distances between X-linked residues in each structural model (Fig. 3). As can be seen in Table 2, 100% of distances in each case are compliant with the accepted range for DSS X-linker of <30 Å^30,67^.

**Figure 3.**
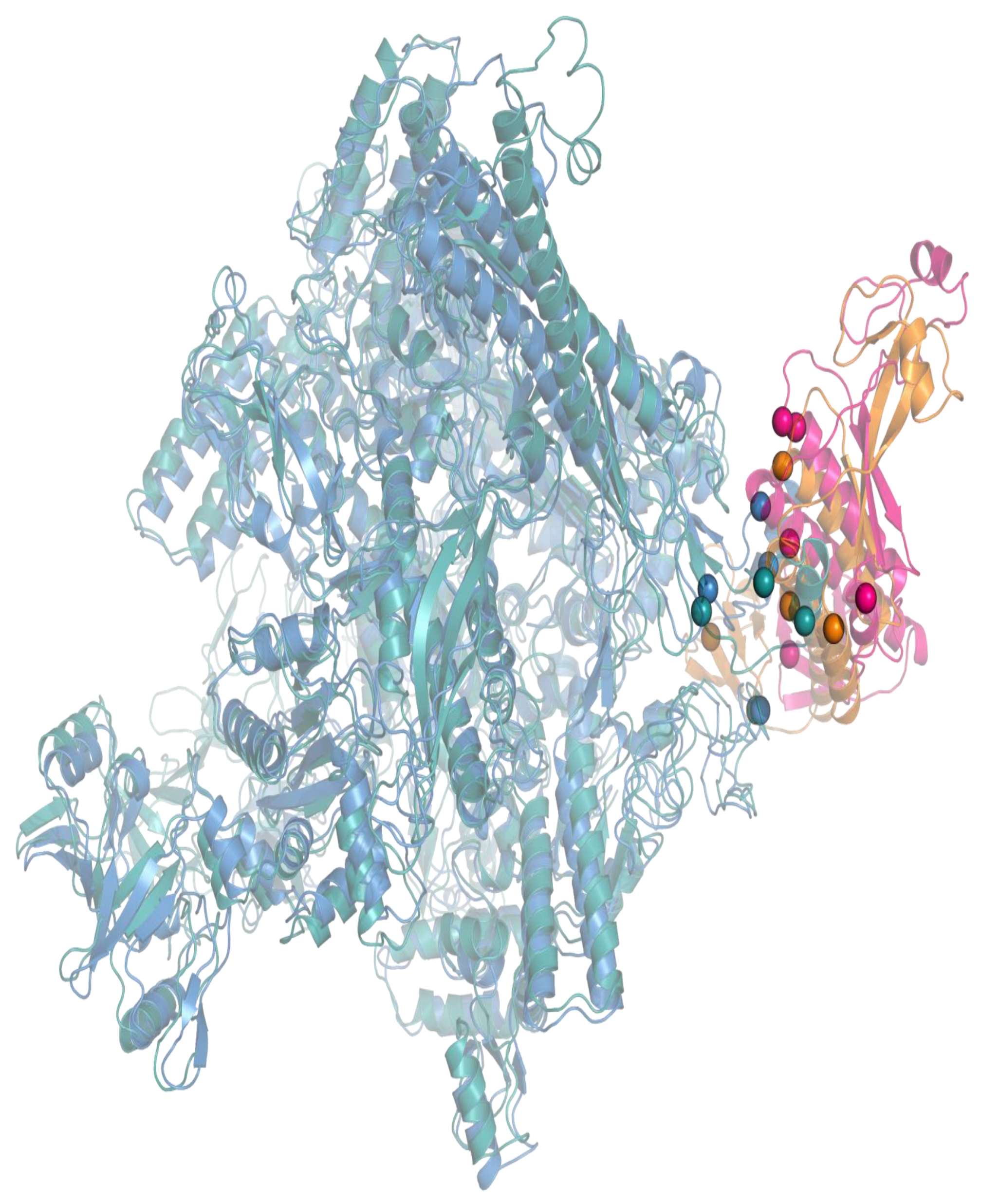
*In vivo* X-linked residues mapped on the docking and cryo EM models of NusA-NTD/RNAP. Structures are pictured as semi-transparent cartoons, CA atoms in X-linked residues – as solid spheres; color scheme is the same as in Fig. 2.

**Table 2.** Euclidean distances between CA-CA atoms in the docking and the experimental (cryo-EM) models of NusA-NTD/RNAP complex

|  | CA-CA distance (Å) |  |
| --- | --- | --- |
|  | cryo-EM | docking |
| NusA3-RpoB900 | 13.9 | 23.6 |
| NusA37-RpoB890 | 26.8 | 18.9 |
| NusA37-RpoB909 | 22.2 | 6.2 |
| NusA38-RpoB890 | 25.5 | 17.4 |
| NusA38-RpoB909 | 20.7 | 6.8 |
| NusA111-RpoB890 | 14.5 | 14.2 |
| NusA111-RpoB909 | 17.8 | 12.5 |
| NusA143-RpoC79 | 12.1 | 18.3 |

Altogether these findings strongly indicate that, although the docking model does not replicate the atomic details of the experimental (cryo EM) one, it sufficiently well reflects the binding poses and the overall architecture of the NusA-NTD/RNAP complex.

## Discussion

We have executed and evaluated the results of an automated workflow for creating a structural model of a protein complex (*E. coli* NusA-NTD/RNAP), which was discovered and structurally interrogated *in vivo* by AP-XL-MS. The X-links-guided docking was performed without any additional input or bias (in fact the docking was completed prior to the time when the reference experimental structural model (6flq) became available). The entire workflow was constructed using freely available, user-friendly computational tools (pLink, HADDOCK, etc), and commercially available reagents. The entire computational pipeline is streamlined, featuring a minimal number of input/output hand-offs compared to other in vivo XL-MS approaches^25,68^.

The efficiency of the workflow reported in this work approaches that of the top-performing DSSO-based methodology^25^, while being more accessible to the laboratories lacking expertise necessary for its implementation. Utilizing broadly cell-permeable X-linker DSS, this workflow outperforms *in vivo* XL-MS approach based on the functionalized X-linker bis(succinimidyl)-3-azidomethyl-glutarate (BAMG)^68^. Unlike DSS, BAMG is not readily available from commercial sources, exhibits restricted cell permeability (e.g. ineffective in treatment of *E. coli*) and toxicity, and, despite functionalization towards X-link enrichment and robust discovery^69^, delivers low X-link discovery rates (84 inter-protein X-links from the entire *B. subtilis* proteome (4260 proteins))^68^.

Targeted AP-XL-MS reported in this work in addition to the structural interrogation of the cell interactome during rapid growth of bacteria can be also applied to mammalian cells cultures, and the discovery of the conditional changes in PPC structure and composition in different physiological conditions (stationary phase, onset of virulence, heat shock, etc), or as response to administration of drugs and other bioactive compounds.

## Materials and Methods

### Buffer components and consumables

Proteomics-grade DSS and DMSO were acquired from Proteochem. Buffer components for cross-linking and protein purification were BioUltra grade (Millipore Sigma). LC-MS/MS was carried out with Thermo Scientific LC-MS grade reagents and solvents. Growth media components were from Thermo Fisher, protease inhibitor cocktail (ProBlock Gold Bacterial 2D) was from Gold Biotechnology, Ready-Lyse lysozyme solution was from Epicentre Biotechnologies, His-Mag Sepharose Ni – from GE. Low-binding pipet tips (Corning DeckWorks) and tubes (Protein LoBind, Eppendorf) were used throughout the experimental workflow.

### *In vivo* X-linking and affinity purification of the RpoC-baited PPCs

*E. coli* strain RL721 (*rpoC*::His6) (generously contributed by Robert Landick, University of Wisconsin-Madison) was grown with agitation at 37^°^C in 0.5X phosphate-buffered Terrific Broth^70^. DSS was dissolved in anhydrous DMSO (300 mM stock) and added to the bacterial culture at OD_600_=0.5 at the final concentration of 2 mM. After 30 min incubation X-linking was quenched by the addition of Tris base to the final concentration of 50 mM. Cells were harvested by centrifugation (4^°^C, 10 min, 6 000 g), resuspended in lysis buffer (50 mM HEPES (pH 7.5), 500 mM NaCl, 1X ProBlock Gold Bacterial 2D) and lysed by the combined action of ultrasonication and Ready-Lyse lysozyme (30 KU/ml). Extract was cleared by centrifugation (4^°^C, 2X 30 min, 29 500 g) and combined with His-Mag Sepharose Ni, the mix was incubated at 4^°^C for 8 hrs with rotation. Unbound material was removed using magnetic separation, the beads were washed with denaturing buffer (8 M urea, 50 mM HEPES (pH 7.5), 500 mM NaCl) at 4^°^C for 2 hrs with rotation, twice with non-denaturing wash buffer (50 mM HEPES (pH 7.5), 500 mM NaCl) at 4^°^C for 2 hrs with rotation, the proteins were eluted in non-denaturing wash buffer, supplemented with 400 mM imidazole.

**Processing of X-linked samples for and by LC-MS/MS** was carried out by NYU Langone Health Proteomics Laboratory as described before^71^.

### Discovery of the *in vivo* X-links

Discovery of the DSS X-links was carried out using pLink1^36^. X-linked peptide search space was defined by combining protein sequences discovered in the enumerative analysis into a single *fasta* file. Search parameters were defined in the *pLink.ini* file by setting the enzyme name to trypsin, maximal number of missed cleavages to 3, maximal e-value to 0.001. Amino acid modifications were limited to 1 constant (Carbamidomethyl[C]), and 3 variable (Oxidation_M, Gln->pyro-Glu, and N-acetyl_Protein) ones. Example of the *pLink.ini* files is included in the Supplement.

### XL-MS-guided protein-protein docking and analysis of the docking models

Distance restraints-guided protein-protein docking was carried out using the Expert interface of the HADDOCK server^59^. Distance restraints were recorded in the *unambig.tbl* file. Starting structures and the *unambig.tbl* files are included in the Supplement. Homology modeling was carried out using the I-TASSER server^56^. Starting structures refinement was carried our using FG-MD^57^ and YASARA^72^ energy minimization servers, docking model refinement was performed using Refinement interface of the HADDOCK server. Coordinate files (*pdb*) manipulations (chain joining and renumbering) was carried out using YASARA Dynamics^73^. Comparison between the top docking model and the reference experimental structure was carried out using TM-score server^66^.

## Acknowledgements

This work was supported by the NIH grant R01 GM107329 and by the Howard Hughes Medical Institute.

## Contributions

EN and VS conceived the study. VS carried out the experiments and data analysis under EN supervision. EN and VS wrote the manuscript.

## Materials and Correspondence

All correspondence and requests for materials should be submitted to Evgeny Nudler. Technical questions may also be directed to Vladimir Svetlov.

